# Ketosis rescues frataxin deficiency and corrects disease phenotypes in an FRDA animal model

**DOI:** 10.64898/2026.01.14.699352

**Authors:** Yina Dong, Miniat W. Adeshina, Ryan Smith, Lucie Vanessa Ngaba, Jacob An, Olivia Seline, Jennifer Coulman, Shahd AL Haj Ali, Clementina Mesaros, Peining Xu, David R Lynch

## Abstract

Friedreich ataxia (FRDA) is an autosomal recessive neurodegenerative disease caused by deficiency of the mitochondrial protein frataxin. Effective therapeutic options remain limited for FRDA. We previously demonstrated that frataxin regulates ketone body metabolism by modulating 3-Oxoacid CoA-Transferase 1 (OXCT1), the rate-limiting enzyme in ketone body catabolism. However, the mechanisms governing frataxin-dependent control of OXCT1 turnover as well as the contribution of frataxin deficiency-induced OXCT1 reduction to FRDA pathogenesis, have remained unclear. Here, we demonstrate that frataxin regulates OXCT1 protein turnover by inhibiting its ubiquitination and subsequent proteasomal degradation. The N-terminal 40 amino acids of frataxin mediate such events, as overexpression of this region alone blocks ubiquitin-proteasome system (UPS)-dependent OXCT1 degradation. To evaluate the impact of OXCT1 deficiency on FRDA phenotypes, we enhanced OXCT1 reduction by introducing a 50% *OXCT1* knockout into frataxin-deficient KIKO mice (KIKO/*OXCT1*⁺/⁻). While OXCT1 deficiency potentiates cell death *in vitro* in control and FRDA patient fibroblasts, further OXCT1 reduction in KIKO mice induces ketosis, increases frataxin levels, and improves neurobehavioral performance. The increase in frataxin does not reflect elevated *FXN* gene transcription but rather enhanced mitochondrial biogenesis, evidenced by increased biogenesis markers, restored mitochondrial morphology and size, and increased mitochondrial gene expression. Fasting-which promotes ketosis-similarly increases frataxin levels and mitochondrial biogenesis markers in older KIKO/*OXCT1*+/− mice. β-hydroxybutyrate administration in FRDA iPSC-derived cardiomyocytes elevates frataxin levels and mitochondrial biogenesis markers, further supporting the beneficial effect of ketosis on frataxin expression and mitochondrial biogenesis. Collectively, our findings demonstrate that ketosis partially restores frataxin levels and ameliorates FRDA-related phenotypes, providing a potential therapeutic strategy for FRDA.

## 1. Introduction

Friedreich ataxia (FRDA) is an autosome recessive neurodegenerative disease characterized by progressive gait and limb ataxia, hypertrophic cardiomyopathy, dysarthria, scoliosis, and in some individuals diabetes (1). FRDA is caused in 94% of the cases by homozygous expanded guanine–adenine–adenine (GAA) repeats in the first intron of *FXN*, with a small percentage (4%) being caused by compound heterozygosity for a non -GAA mutation in one allele and an expanded GAA repeat on the other (1). The expanded GAA repeats silence *FXN* transcription and consequently reduce the levels of the frataxin protein product (2). A lack of frataxin compromises energy and calcium metabolism (3–7), disrupts maintenance of mitochondrial membrane potential (5, 8), and leads to iron accumulation and oxidative stress (9–10). These factors contribute to mitochondrial failure leading to the pathological changes in tissues with a high energy demand.

3-Oxoacid CoA-Transferase 1 (OXCT1), the rate limiting enzyme in ketone body catabolism, catalyzes the conversion of ketone bodies to acetoacetyl-CoA that is then fed into the Krebs cycle for ATP production. Ketone bodies provide an alternative energy source to glucose for heart, brain, and skeletal muscle during times of low glucose levels such as fasting and prolonged exercise. Germline OXCT1 knockout (KO) mice, unable to oxidize ketone bodies in any tissue, die within 48 hours after birth due to hyperketonemic hypoglycemia (11). In humans, mutations in the *OXCT1* gene cause OXCT1 deficiency, leading to recurrent episodes of ketoacidosis, a life-threatening metabolic disorder (12–14). Moreover, conditional knockout of *OXCT1* in peripheral sensory neurons results in severe proprioceptive deficits in mice (15), underscoring the essential role of ketone body metabolism in sensory nervous system development and function.

Beyond their role as alternative energy substrates, ketone bodies also function as signaling molecules that exert epigenetic and post-translational effects, promoting improved mitochondrial function, enhanced antioxidant capacity, and anti-inflammatory responses (16–19). More recently, β-hydroxybutyrate (BHB), the predominant circulating ketone body, regulates gene expression through covalent modification of histone lysine residues via β-hydroxybutyrylation (Kbhb) (20). These findings link ketosis to covalent chromatin modifications and expand the role of ketone bodies beyond fuel provision to that of dynamic regulators of chromatin architecture and gene transcription. Ketosis-based therapies, such as the ketogenic diet, have been used in neurological disorders including epilepsy, with proposed mechanisms involving enhanced mitochondrial biogenesis and function (21). The impact of ketosis on FRDA has not been systematically investigated.

We previously demonstrated that direct interaction with frataxin stabilizes OXCT1, protecting it from ubiquitin–proteasome system (UPS)-mediated degradation (22). Frataxin deficiency reduces OXCT1 protein levels across multiple tissues, including those from FRDA patients, resulting in impaired ketone body utilization and elevated circulating ketone levels (ketosis) (22). However, the pathological and potentially adaptive consequences of OXCT1 deficiency–induced disruption of ketone body metabolism in FRDA remain unclear. In the present study, we identify a beneficial effect of OXCT1 deficiency–induced ketosis on frataxin levels and neurobehavioral outcomes in FRDA, mediated in part through enhanced mitochondrial biogenesis.

## 2. Materials and Methods

### 2.1 Animals

C57BL/6 mice (stock no: 000664) and frataxin knock-in/knockout (KIKO) mice (B6.Cg-Fxntm1.1Pand Fxntm1Mkn/J; stock no. 012329) were purchased from Jackson Laboratory. C57BL/*6-OXCT1tm1a*(KOMP)Wtsi/Mmucd mice were purchased from the Mutant Mouse Resource & Research Centers (MMRRC) repository (University of California, Davis). *KIKO/OXCT1+/−* mice were generated through a series of crossing among KOWT, KIKI, and *OXCT1+/−* mice. Mice handling and treatment were in accordance with standard regulations approved by the Children’s Hospital of Philadelphia Institutional Animal Care and Use Committee (IACUC; protocol 16–250).

### 2.2 Accelerating rotarod

Rotarod test was performed as described previously (23).

### 2.3 Preparation of tissue homogenates

After mice were euthanized, the cerebellum and skeletal muscle were collected and homogenized as described previously (24). The homogenization buffer contained 100 mM Tris-HCl (pH 7.4), 150 mM NaCl, 1% IGEPAL, 1 mM EDTA, and 0.5% sodium deoxycholate, supplemented with a protease inhibitor cocktail (1:500 dilution; Calbiochem, Darmstadt, Germany). Homogenates were incubated at 4 °C for one hour, then centrifuged at 13,000 rpm for 15 minutes. The resulting supernatant was stored at −80 °C until use.

### 2.4 Overexpression of plasmid DNAs in HEK293 cells

Overexpression of plasmid DNAs is performed as previously (25). HEK293 cells were transfected with plasmid DNAs containing Cyto-HA-Ubiquitin (Addgene, Watertown, MA), Mito-HA-Ubiquitin (A generous gift from Dr. Giovanni Bénard, Université de Bordeaux, France), wildtype (WT) frataxin or frataxin truncation mutants (1–190, 1–168, and 1–141 aa) fused to a C-terminal HA tag (Genscript, Piscataway, NJ), WT frataxin or frataxin truncation mutants (1-120, 1-100, 1-80, 1-55, 1-40 aa) fused to a C-terminal GFP tag (Genscript, Piscataway, NJ) using Lipofectamine™ 2000 reagent (Thermo Fisher Scientific Inc.) for 24 h. Cells were then collected and subjected to co-immunoprecipitation or western blot.

### 2.5 Total DNA and RNA isolation and quantification

As previously described (22, 26), total DNA was extracted from the cerebellum of control, KIKO and KIKO/*OXCT1+/−* mice at 5-7 M of age using DNeasy blood and tissue kit (Qiagen, Germantown, MD). Total RNA was extracted from the cerebellum of control, KIKO and KIKO/*OXCT1*+/− mice at 5-7 M of age using Qiagen’s RNeasy Mini kit (Qiagen, Germantown, MD). RNA was then reverse-transcribed into cDNA using Bio-Rad’s iScripttM cDNA Synthesis Kit (Bio-Rad, Philadelphia, PA). Both DNA and RNA were quantified by a NanoDrop 2000 Spectrophotometer (Thermo Fisher Scientific, Waltham, MA).

### 2.6 Quantitative polymerase chain reaction (qPCR)

qPCR was performed using SYBR kit (Qiagen, Hilden, Germany) in Applied Biosystem Real Time PCR instrument (Thermo Fisher Scientific, Waltham, MA). The primer sequences for mt-ND1, Cftr, frataxin and actin were used as previously described (22). Primers were synthesized by Integrated DNA Technologies, Inc. (Coralville, IA), and DNA or gene transcript levels were normalized to the expression levels of beta actin.

### 2.7 Induced pluripotent stem cell (iPSC) culture and cardiac differentiation

#### 2.7.1 iPSC Culture and maintenance

iPSCs were plated on a human embryonic stem cell (hESC)-Qualified Matrix Matrigel (Corning, Corning, NY, 354277) coated 60mm dish. Thawing media consisted of mTeSR Plus (STEMCELL Technologies, Vancouver, BC, 100-0276) and 10uM of Rock Inhibitor (STEMCELL Technologies, Vancouver, BC, Y27632) at 1:1000 dilution. After 24 hours, media was changed to mTeSR Plus alone. Cells were maintained in mTeSR Plus up until reaching 70-80% confluency, determined by the size of the colonies. Once confluent, Dispase (STEMCELL Technologies, Vancouver, BC, 07923) was used to obtain a single cell suspension. Cells were then counted and plated in 6 well plates. Cells were seeded at several different densities (2×10^5^ to 1×10^6^ cells/well), as optimal seeding density for cardiac differentiation was cell line dependent.

#### 2.7.2 Cardiac Differentiation

Cardiac differentiation was performed using a previously published small-molecule protocol based on initial inhibition of GSK3 signaling followed by WNT signaling inhibition (27–28). Cells were maintained in standard conditions (mTeSR Plus) until reaching 85-95% confluence (day x – day –1). At d0, media was changed to *Differentiation Medium* (500 ml RPMI 1640 with HEPES with GlutaMax (Thermo Fisher Scientific, Waltham, MA, 72400047), 250 mg human recombinant Albumin (Sigma-Aldrich, St. Louis, MO, A9731-10G), and 100 mg L-ascorbic Acid 2-Phosphate (Sigma-Aldrich, St. Louis, MO, A8960-5G)) supplemented with 4µM of CHIR99021 (Tocris, 4423). At d2, media was replaced with *Differentiation Medium* supplemented with 5uM IWP2 (Tocris, Bristol, UK, 3533). On d4-d6, cells were maintained in *Differentiation Medium* without any supplementation. Starting at d8, media was replaced with RPMI 1640 with 2% B27 Supplement with Insulin (Thermo Fisher Scientific, Waltham, MA, 17504044, 50x). The first spontaneously beating cells typically appeared on d8 or d9. Cardiac selection was performed between d12 and d40 of differentiation due to the highly proliferative state (high sensitivity to lactate) by replacing media with *Differentiation Medium* supplemented with 2mL of 1M Lactate/HEPES (Thermo Fisher Scientific, Waltham, MA, 15630080) for 4-5 days. After this, media was replaced and maintained in RPMI 1640 with 2% B27 Supplement with Insulin.

### 2.8 Western blot

Western blot was performed as described previously (24)). The following antibodies were used: OXCT1 (Thermo Fisher Scientific Inc., Waltham, MA, 1/1000), frataxin (Abcam, Cambridge, United Kingdom, 1/500), pan-actin (Cell signaling, Danvers, MA, 1/1000), Tfam (Abcam, Cambridge, United Kingdom, 1/1000), Nrf2 (Abcam, Cambridge, United Kingdom, 1/1000), HA (Cell signaling, Danvers, MA, 1/1000), GFP (Proteintech Group, Rosemont, IL, 1/1000), GRP75 (Abcam, Cambridge, United Kingdom, 1/1000), histone 3 (Cell signaling, Danvers, MA, 1/1000), acetylated histone 3 lysine 9 (H3K9) (Cell signaling, Danvers, MA, 1/1000) and acetylated histone 3 lysine 14 (H3K14) (Thermo Fisher Scientific, Waltham, MA).

### 2.9 Co-Immunoprecipitation

Co-Immunoprecipitation was performed as described previously (24). The following antibodies are used: HA (Cell signaling, Danvers, MA), OXCT1 (Proteintech, Rosemont, IL), GFP (Proteintech Group, Rosemont, IL).

### 2.10 Protein stability assay

HEK293 cells were cultured as previously described (25). Cells were transfected for 24 h with either vector control cDNA or frataxin 1–40 bearing a C-terminal HA tag, then treated with cycloheximide (50 μg/ml, Sigma, St. Louis, MO) or vehicle for varying durations (0, 1, 2, 3, 4, and 6 h). Following treatment, cells were lysed for Western blot analysis.

### 2.11 Immunofluorescence

Immunofluorescence was performed as previously described (25). Mitotracker^TM^ Red CMXRos (Thermo Fisher Scientific, Waltham, MA) was loaded to HEK293 cells for 45 minutes before immunofluorescence was performed. The following antibodies were used: HA (Cell Signaling, Danvers, MA), OXCT1 (Thermo Fisher Scientific Inc., Waltham, MA), GFP (Neuromab, Davis, CA at 1/200).

### 2.12 Transmission electron microscopy

For the ultrastructural studies, control, KIKO and KIKO/*OXCT1*+/− mice of either sex at 5-7 M (n=3 per group) were perfused with 2.5% glutaraldehyde and 2% paraformaldehyde, after which their cerebella were collected. Samples for electron microscopy were fixed overnight at 4 °C in 2.5% glutaraldehyde and 2.0% paraformaldehyde in 0.1 M sodium cacodylate buffer (pH 7.4), then processed and stained at the Electron Microscopy Resource Laboratory, University of Pennsylvania Perelman School of Medicine Biomedical Research Core Facilities.

### 2.13 LC-MS analysis

LC-MS was performed as described previously (22). Blood collected from control, KIKO, and KIKO/*OXCT1+/−* mice was centrifuged at 15,000 rpm for 15 min, and the resulting serum was collected for BHB measurement.

### 2.14 Statistical analysis

Statistical differences between two groups were analyzed using a two-tailed Student’s t-test, and comparisons among three or more groups were evaluated using one-way ANOVA followed by Bonferroni’s post hoc test. A p-value of less than 0.05 was considered statistically significant.

## 3. Results

### 3.1 Frataxin prevents proteasomal degradation of OXCT1 by inhibiting its ubiquitination

We previously showed that frataxin regulates OXCT1 by preventing its UPS-dependent degradation (22). Because proteasomal degradation can occur with or without ubiquitination, we next examined whether OXCT1 is ubiquitinated prior to its degradation. HEK293 cells transfected with Cyto-HA-ubiquitin or vector control were treated with MG132 or vehicle for 5 hours, followed by OXCT1 immunoprecipitation. Overexpression of Cyto-HA-ubiquitin increased the level of ubiquitinated OXCT1 relative to vector control, and MG132 treatment further enhanced cytosolic ubiquitinated OXCT1 compared with vehicle treatment (**Figure 1A**), indicating that OXCT1 is subject to ubiquitination.

**Figure 1.**
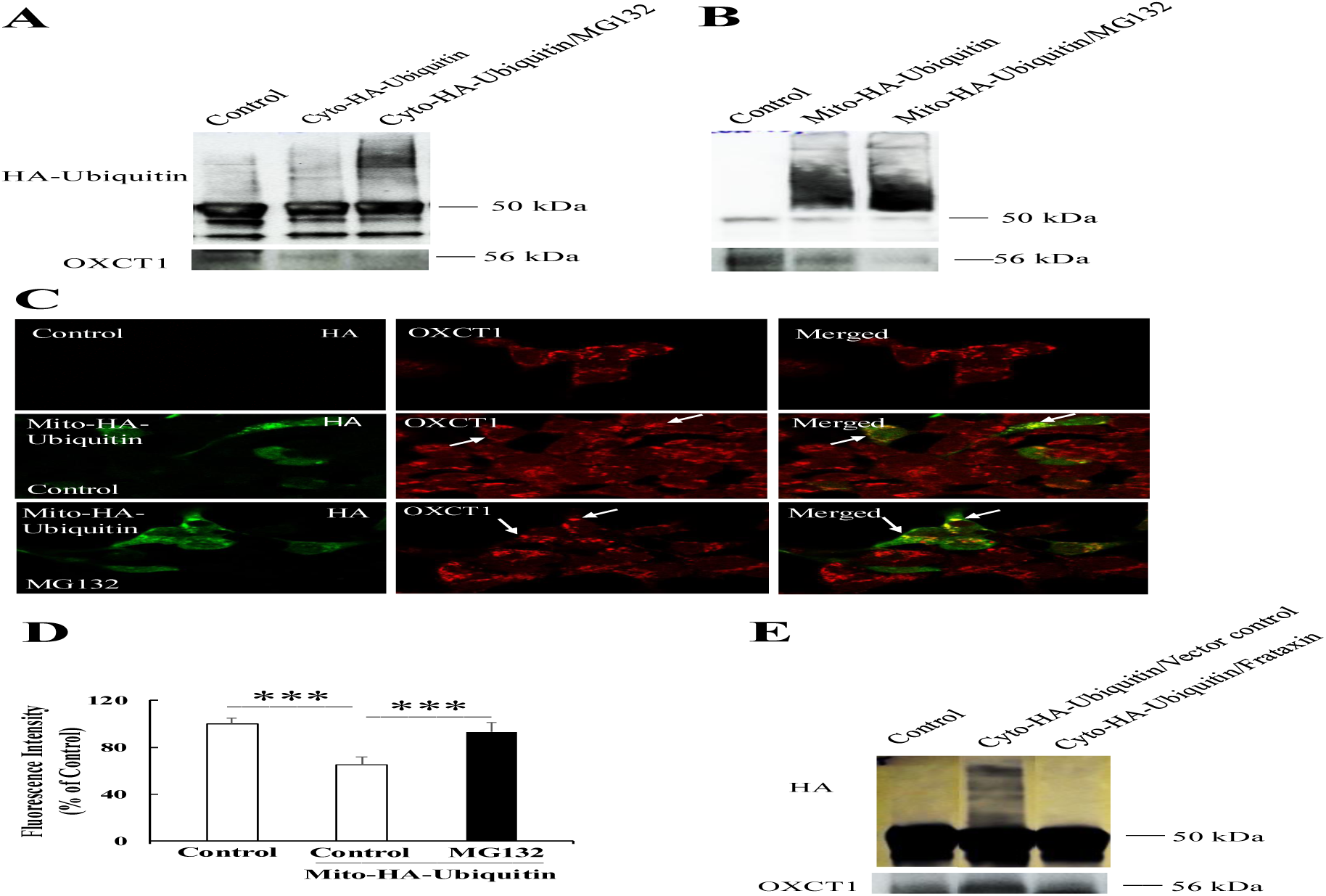
Frataxin inhibits the ubiquitination of OXCT1. HEK293 cells transfected with vector control, Cyto-HA-ubiquitin, or Mito-HA-ubiquitin were treated with MG132 (10 μM) or vehicle control for 5 hours, followed by immunoprecipitation with an anti-OXCT1 antibody and Western blot analysis. Representative blots show ubiquitinated OXCT1 in both the cytosolic (A) and mitochondrial (B) fractions, with MG132 treatment markedly enriching ubiquitinated OXCT1 in both compartments. In HEK293 cells transfected with Mito-HA-ubiquitin, OXCT1 fluorescence intensity was significantly reduced compared with vector control-transfected controls, whereas MG132 treatment restored OXCT1 fluorescence intensity (C, D; n = 30 cells, \*\*\**P* < 0.001). Co-transfection with frataxin markedly reduced OXCT1 ubiquitination in the cytosolic fraction (E), indicating that frataxin regulates OXCT1 protein turnover by blocking its ubiquitination. Data were expressed as mean ± SE (error bars).

Because OXCT1 is a mitochondrial protein, we next evaluated whether its ubiquitination also occurs within mitochondria. HEK293 cells transfected with mitochondria-targeted HA-ubiquitin (Mito-HA-ubiquitin) or vector control were treated with MG132 or vehicle for 5 hours, followed by OXCT1 immunoprecipitation. No ubiquitinated OXCT1 was detected in cells transfected with vector control, whereas overexpression of Mito-HA-ubiquitin significantly increased ubiquitinated OXCT1. MG132 treatment further enriched mitochondrial ubiquitinated OXCT1 (**Figure 1B**), consistent with inhibition of ubiquitination-dependent OXCT1 degradation. Supporting these findings, immunofluorescence revealed colocalization of OXCT1 with Mito-HA-ubiquitin (**Figure 1C**). Mito-HA-ubiquitin overexpression also reduced OXCT1 fluorescence intensity, while MG132 treatment restored and enriched OXCT1 staining (**Figure 1C and 1D)** (Control/Mito-HA-Ubiquitin: 35% decrease from vector control, n=30 cells, *P*<0.001; MG132/Mito-HA-Ubiquitin: 28% increase from vehicle control, n=30 cells, *P*<0.001), further indicating that OXCT1 degradation is ubiquitination-dependent. These results also demonstrate that OXCT1 ubiquitination can occur in mitochondria.

Because frataxin inhibits the proteasomal degradation of OXCT1 in a ubiquitination-dependent manner (**Figure 1A-1D**), we next examined whether frataxin interferes with OXCT1 ubiquitination. HEK293 cells were transfected with Cyto-HA-ubiquitin with or without frataxin overexpression, followed by OXCT1 immunoprecipitation. The presence of frataxin markedly reduced OXCT1 ubiquitination (**Figure 1E**), indicating that frataxin prevents OXCT1 degradation by blocking its ubiquitination.

### 3.2 Frataxin interacts with OXCT1 through its N-terminal 1-40 amino acids

To identify the region required for OXCT1 interaction, we generated a series of C-terminal frataxin deletion mutants (**Figure 2A**). HA-tagged WT or frataxin truncations (1–190 aa, 1–168 aa, and 1–141 aa) were expressed in HEK293 cells for 24 hours, followed by co-immunoprecipitation assays. HA but not IgG immunoprecipitation pulled down OXCT1 in cells expressing WT frataxin, and all three truncation mutants displayed comparable binding to OXCT1 (**Figure 2B**). These results indicate that the C-terminal 142–210 amino acids of frataxin are dispensable for its interaction with OXCT1.

**Figure 2.**
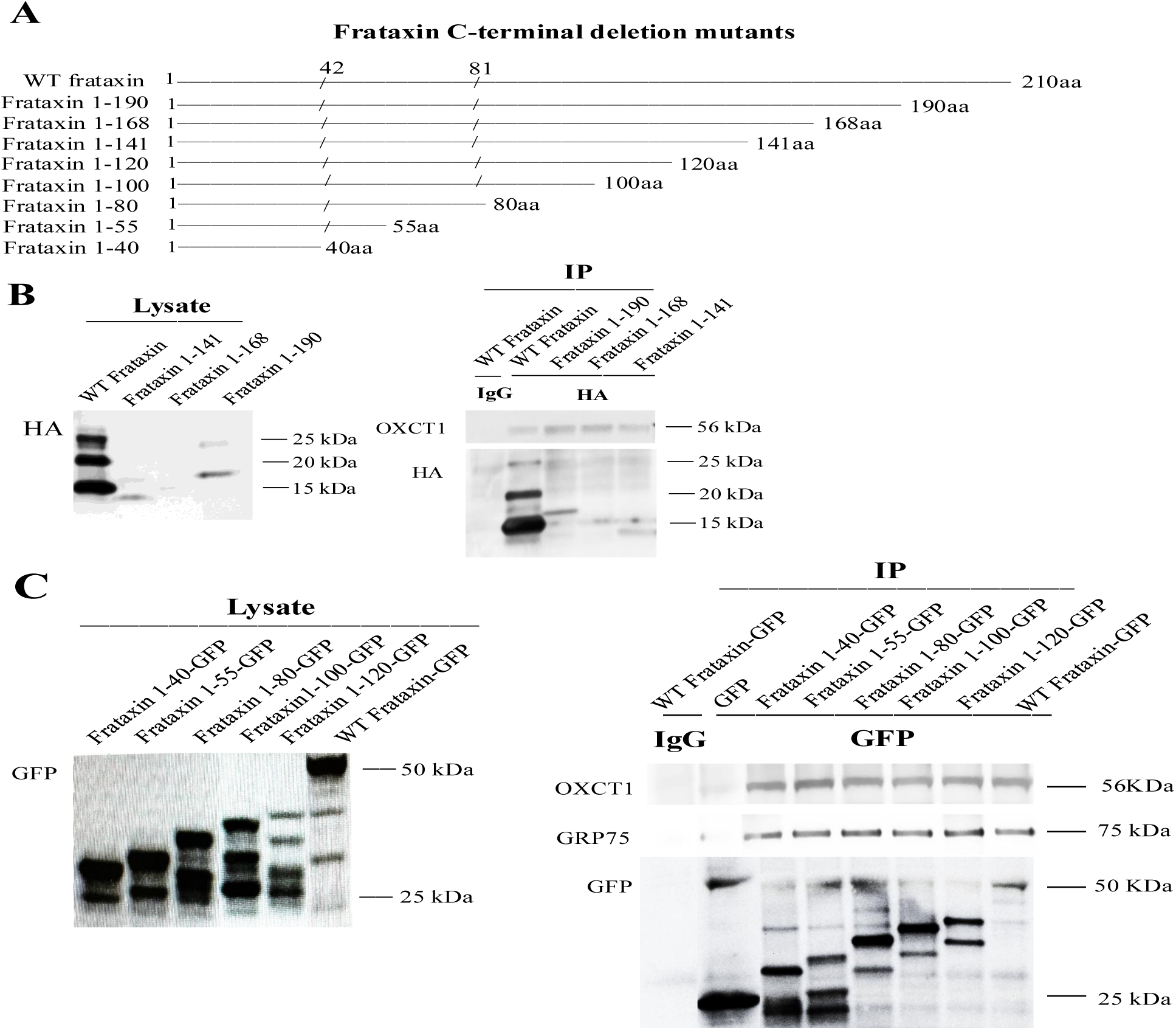
Frataxin binds OXCT1 through its N-terminal 40 amino acids. (A) Schematic representation of C-terminal truncation mutants of the frataxin protein. (B) Immunoprecipitation with an anti-HA antibody revealed comparable levels of OXCT1 co-precipitated from HEK293 cells expressing frataxin 1–190-HA, 1–168-HA, or 1–141-HA constructs relative to WT frataxin. IgG was used as a negative control. (C) Similar levels of OXCT1 were also immunoprecipitated from HEK293 cells expressing N-terminal frataxin fragments (1–120-GFP, 1–100-GFP, 1–80-GFP, 1–55-GFP, and 1–40-GFP) compared with WT frataxin. GRP75 and GFP served as positive and negative controls, respectively.

We next examined which region within amino acids 1-141 is required for OXCT1 binding. GFP-tagged frataxin truncations (1-120 aa, 1-100 aa, 1-80 aa, 1-55 aa, 1-40 aa) along with WT frataxin were expressed in HEK293 cells and subjected to co-immunoprecipitation. GFP, but not IgG, immunoprecipitation pulled down OXCT1 in cells expressing WT frataxin, and all five truncation mutants retained similar binding to OXCT1 (**Figure 2C**). Likewise, all five truncation mutants bound GRP75, a known frataxin chaperone (25). GFP-only transfected cells served as a negative control, and no OXCT1 was detected in GFP immunoprecipitates (**Figure 2C**). The comparable binding of frataxin 1–40 to OXCT1 to that of WT frataxin indicates that amino acids 41–210 are dispensable for frataxin–OXCT1 interactions, and that the OXCT1-binding domain of frataxin resides within its N-terminal 1–40 amino acids.

To determine whether frataxin residues 1–40 have a function similar to that of WT frataxin in regulating OXCT1 protein turnover, we transfected HEK293 cells with either a vector control or a plasmid encoding frataxin 1–40 with a C-terminal HA tag. OXCT1 turnover was then examined in the presence of cycloheximide, a protein synthesis inhibitor. Treatment with cycloheximide for 2 h decreased OXCT1 protein levels (**Figure 3A and B) (**29% decrease, n = 6, *P* < 0.05). This decrease continued until 6 h after cycloheximide treatment (**Figure 3A and B) (**32%, 46%, and 56% decrease for 3, 4, and 6 h, respectively, n = 6, \**P* < 0.05, \*\**P* < 0.01). Frataxin 1–40-HA overexpression prevented OXCT1 degradation, with levels remaining stable across multiple time points, indicating that the 1–40 amino acid fragment of frataxin is functional.

**Figure 3.**
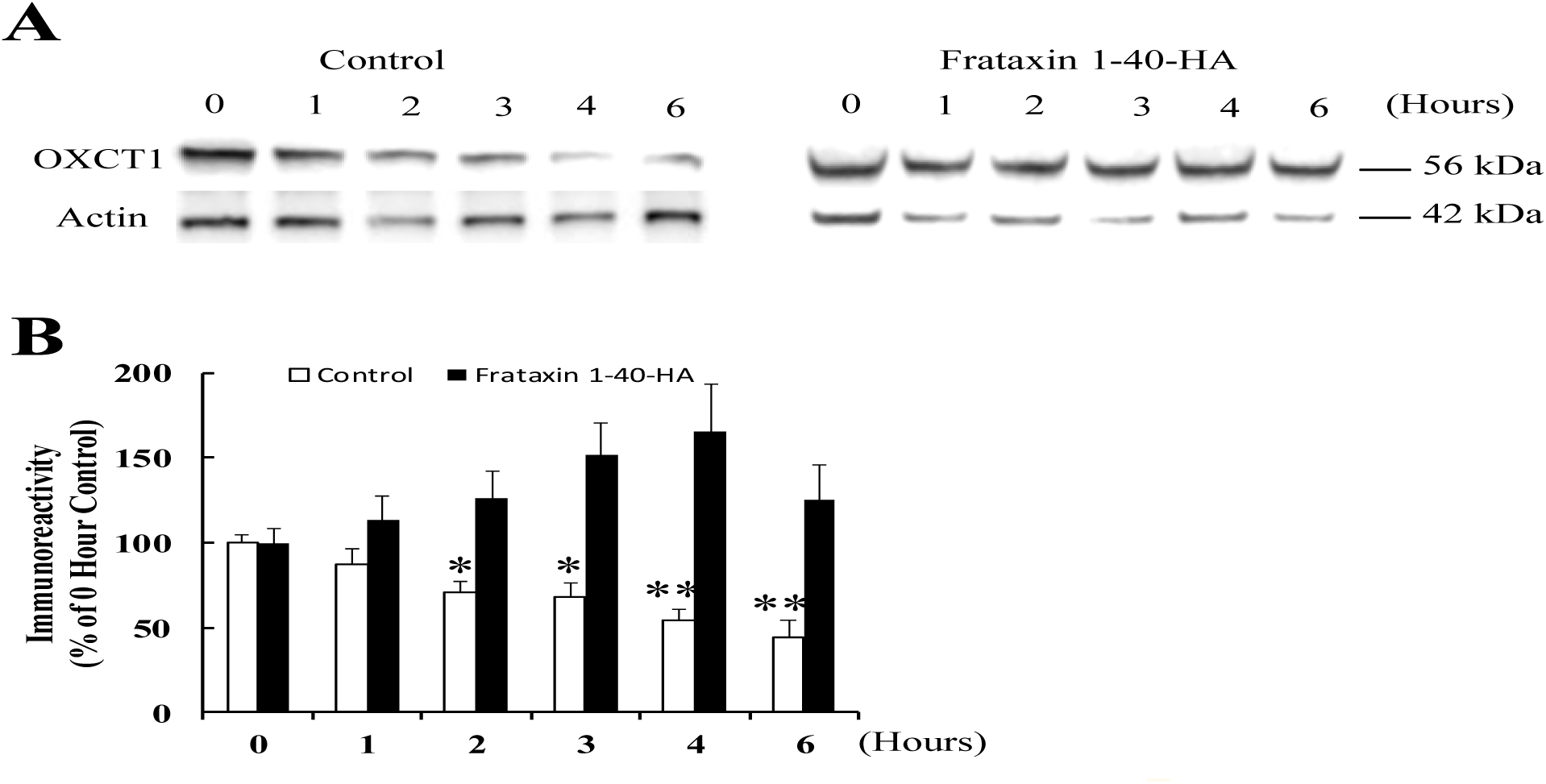
Frataxin 1-40 blocks UPS-mediated OXCT1 degradation. (A) Representative blots showing OXCT1 protein degradation over time in the presence of cycloheximide with or w/o frataxin 1-40-HA overexpression in HEK293 cells. (B) Quantification of OXCT1 degradation in the presence of cycloheximide with or w/o frataxin 1-40-HA overexpression in HEK293 cells. \**P* < 0.05, \*\**P* < 0.01. Data was expressed as mean ± SE (error bars).

### 3.3 OXCT1 reduction potentiates cell death *in* vitro in human skin fibroblasts

We next investigated the functional consequences of OXCT1 deficiency by assessing its impact on FRDA-associated phenotypes. We initially examined how OXCT1 reduction affects cell viability in a cellular model of ferroptosis, a mechanism of cell death proposed in FRDA (29–30), before testing its impact on FRDA abnormalities *in vivo*. Cultured human fibroblasts were subjected to OXCT1 siRNA for 5 days followed by treatment with the ferroptosis inducer erastin for 48 hours and assessment of ATP levels. OXCT1 siRNA transfection efficiently decreased OXCT1 protein levels (**Figure 4A and 4B) (**52% reduction, n=4, *P*<0.05). Erastin treatment had no effect on ATP levels in control fibroblasts but decreased ATP content in FRDA patient-derived fibroblasts (**Figure 4C and 4D) (**43% decrease, n=5, *P*<0.05). OXCT1 knockdown decreased ATP content in both control and FRDA patient fibroblasts (**Figure 4C and 4D) (**Control fibroblasts: 22% decrease, n=4, *P*<0.05; Patient fibroblasts: 66% reduction, n=4, *P*<0.01). These results indicate that OXCT1 deficiency potentiates ferroptotic cell death in both control and FRDA patient-derived fibroblasts.

**Figure 4.**
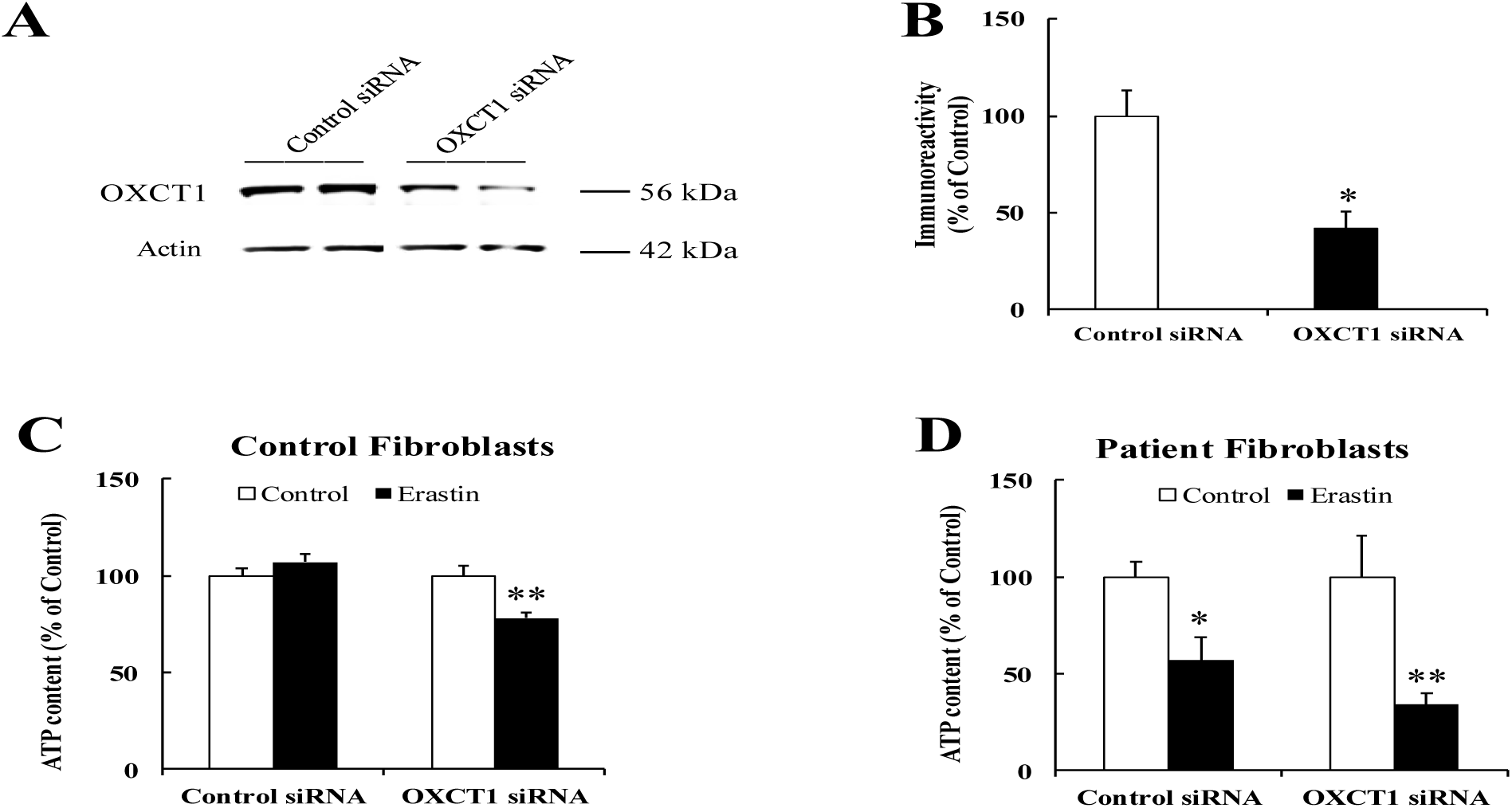
OXCT1 deficiency potentiates cell death in both control and FRDA patient fibroblasts. Representative blots and bar graphs demonstrate decreased OXCT1 protein levels in human control skin fibroblasts transfected with OXCT1 siRNA for 5 days (A and B, n=4). Erastin treatment had no effect on ATP levels in control fibroblasts but significantly decreased ATP content in FRDA patient-derived fibroblasts (C and D, n=5). OXCT1 knockdown decreased ATP content in both control and FRDA patient fibroblasts treated with erastin (C and D, n=4). \**P*<0.05, \*\**P*<0.01. Data was shown as mean ± SE (error bars).

### 3.4 Further OXCT1 deficiency causes ketosis *in vivo* in KIKO/*OXCT1*+/− mice

To study the effect of OXCT1 deficiency on the FRDA phenotypes *in vivo*, we generated the KIKO/*OXCT1*+/− mouse to amplify the OXCT1 deficit and model the effect of mild ketosis on FRDA. The KIKO/*OXCT1*+/− mouse was developed by introducing a 50% knockout of the *OXCT1* gene into the frataxin-deficient KIKO mouse model. KIKO/*OXCT1*+/− mice had lower levels of OXCT1 mRNA and protein in the cerebellum compared to KIKO mice (**Figure 5A, 5B and 5C)** (OXCT1 mRNA: 50% decrease, n=6, *P*<0.001; OXCT1 protein: 46% decrease, n=9, *P*<0.01). Given the large drop in OXCT1 protein levels in KIKO/*OXCT1*+/− mice (69% decrease from control mice, n=9, *P*<0.01), we examined serum BHB levels by mass spectrometry. While heterozygous OXCT1 deficient (*OXCT1*+/−) mice exhibit no change in serum BHB levels (11, 31), KIKO/*OXCT1*+/− mice showed a significant increase in serum BHB levels at physiological state (non-fasting) compared to both control (**Figure 5D) (**35% increase, n=13, *P*<0.05) and KIKO mice (74% increase, n=12, *P*<0.01), suggesting that further OXCT1 reduction impairs ketone body metabolism and causes ketosis in KIKO/*OXCT1*+/− mice at physiological condition.

**Figure 5.**
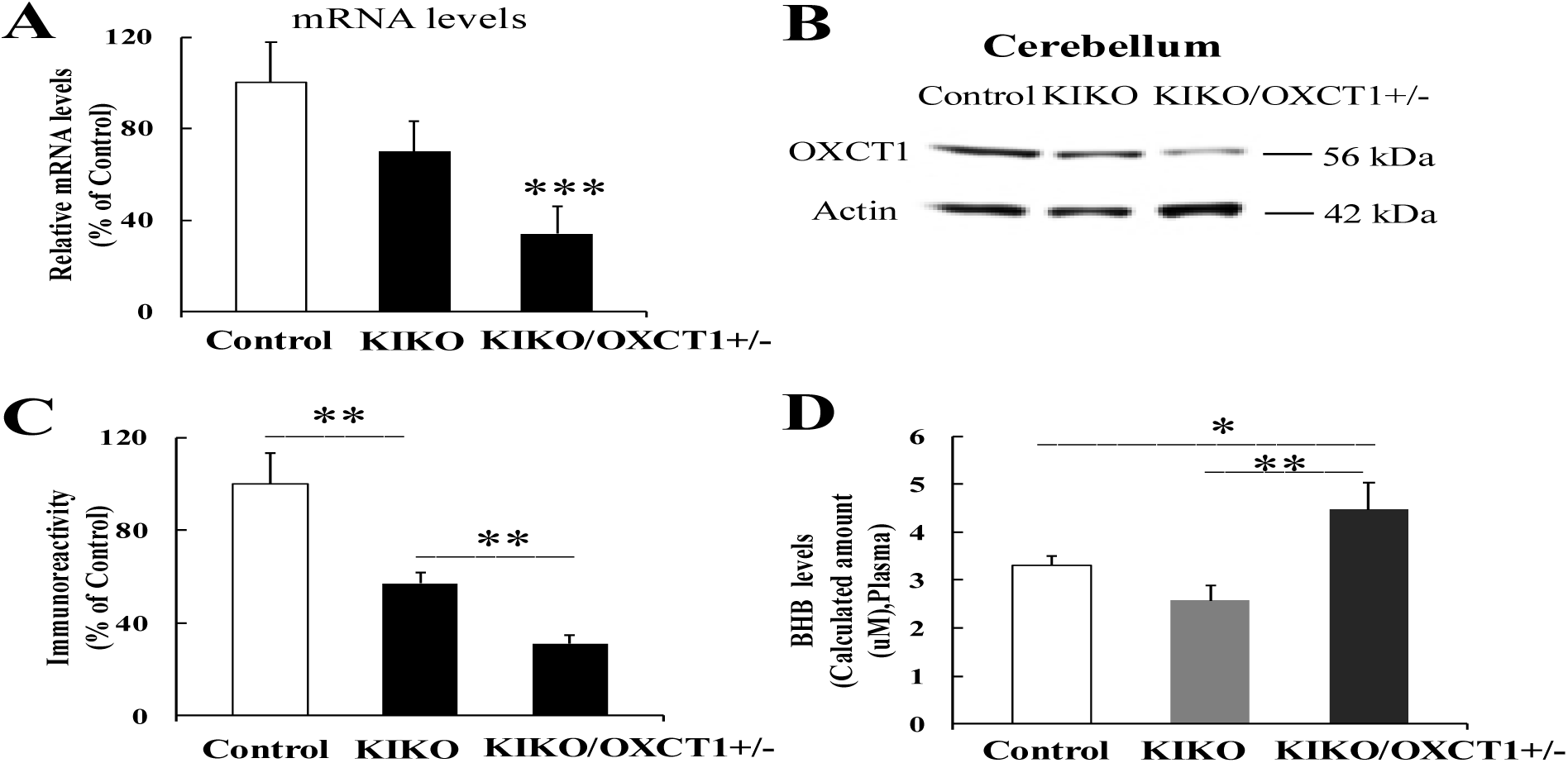
Further OXCT1 reduction causes ketosis in KIKO/*OXCT1*+/− mice. Introduction of a 50% OXCT1 gene knockout into the frataxin-deficient KIKO mouse model resulted in significantly reduced OXCT1 mRNA (A) and protein levels in cerebellar homogenates (B and C, n=8). KIKO/*OXCT1*+/− mice also exhibited significantly elevated serum BHB levels compared with control and KIKO mice (D) (n=13). Data was shown as mean ± SE (error bars). * *p*<0.05, \*\**p*<0.01, \*\*\**p*<0.001.

### 3.5 Further OXCT1 reduction increases frataxin levels and mitochondrial biogenesis markers in the cerebellum of KIKO/*OXCT1*+/− mice

We next examined whether ketosis in KIKO/*OXCT1*+/− mice affects frataxin levels. Compared to KIKO mice at the same age, the cerebellar homogenates of KIKO/*OXCT1*+/− mice (5-7M) had significantly higher levels of frataxin protein (**Figure 6A and 6B**) (2.8-fold increase, n=9, *P*<0.01). As frataxin mRNA levels remained unchanged (**Figure 6E**), we examined if this increase is caused by increased mitochondrial biogenesis. In comparison with KIKO mice, the mitochondrial biogenesis markers Nrf2 and Tfam significantly increased in the cerebellar homogenates of KIKO/*OXCT1*+/− mice (**Figure 6A and 6B)** (41% and 42% increase for Nrf2 and Tfam respectively, n=6, *P*<0.01). In contrast, OXCT1 reduction in control/*OXCT1*+/− mice had no effect on frataxin and the mitochondrial biogenesis marker Tfam **(Figure 6C and 6D) (**n=6, *P*>0.05).

**Figure 6.**
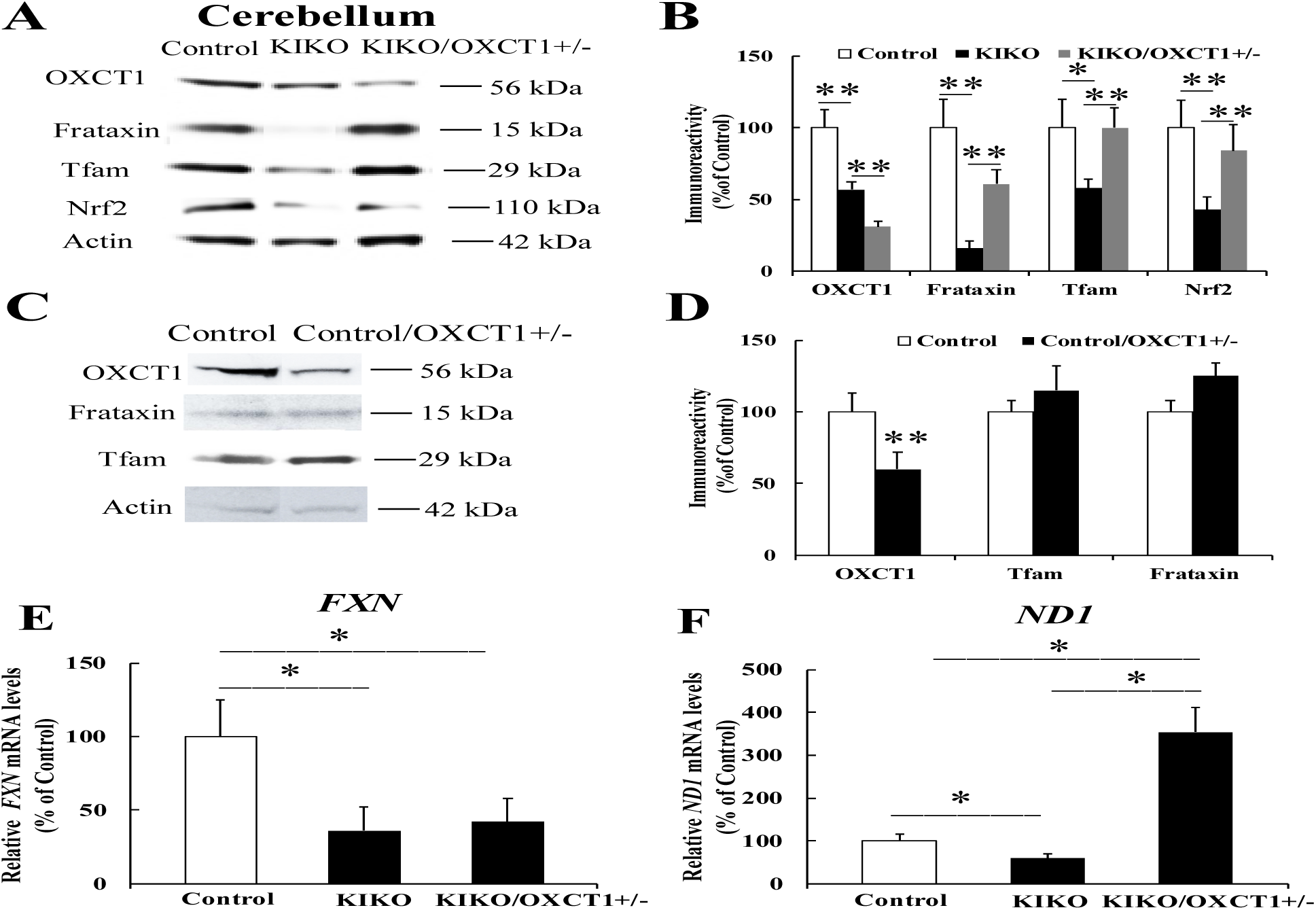
Further OXCT1 reduction increases levels of frataxin and mitochondrial biogenesis markers. Representative blots and bar graphs demonstrate increased levels of frataxin and mitochondrial biogenesis markers in the cerebellar homogenates of KIKO/*OXCT1*+/− mice (A and B, n=9). OXCT1 reduction in control mice had no effect on frataxin and the mitochondrial biogenesis marker Tfam (C and D, n=6). (E) *FXN* mRNA levels in the cerebellum were quantified by RT–qPCR and normalized to actin. (F) *ND1* mRNA levels in the cerebellum were quantified by RT–qPCR and normalized to actin. * *p*<0.05, \*\**p*<0.01. Data was shown as mean ± SE (error bars).

We further confirmed increased mitochondrial biogenesis in KIKO/*OXCT1*+/− mice by measuring mRNA levels of the mitochondria encoded gene NADH dehydrogenase 1 (*ND1)*. Frataxin mRNA levels were markedly reduced in the cerebellum of both KIKO and KIKO/*OXCT1*+/− mice compared with control (**Figure 6E)** (64% and 58% decrease for KIKO and KIKO/*OXCT1*+/− mice respectively, n=5-6, *P*<0.05). In contrast, *ND1* mRNA levels were significantly decreased in KIKO mice compared with control (**Figure 6F**) (41% decrease, n=5–7, *P*<0.05), consistent with impaired mitochondrial biogenesis in these mice (20), but were markedly elevated in KIKO/*OXCT1*+/− mice compared with both KIKO and control mice (**Figure 6F) (**295% and 254% increase for KIKO and control respectively, n=5-6, *P*<0.05). Together, these findings demonstrate that mitochondrial biogenesis is enhanced in KIKO/*OXCT1*+/− mice.

### 3.6 Further OXCT1 reduction improves behavioral phenotypes in KIKO/*OXCT1*+/− mice

Frataxin deficiency induces mitochondrial fragmentation both in cellular and animal models of FRDA (32–34). We next examined whether these mitochondrial morphological abnormalities are rescued in KIKO/*OXCT1*+/− mice. Electron microscopy revealed reduced and fragmented mitochondria in cerebellar Purkinje neurons of KIKO mice. In contrast, both mitochondrial size and morphology were restored to near-normal in cerebellar Purkinje neurons of KIKO/*OXCT1*+/− mice (**Figure 7A**), supporting improved mitochondrial health.

**Figure 7.**
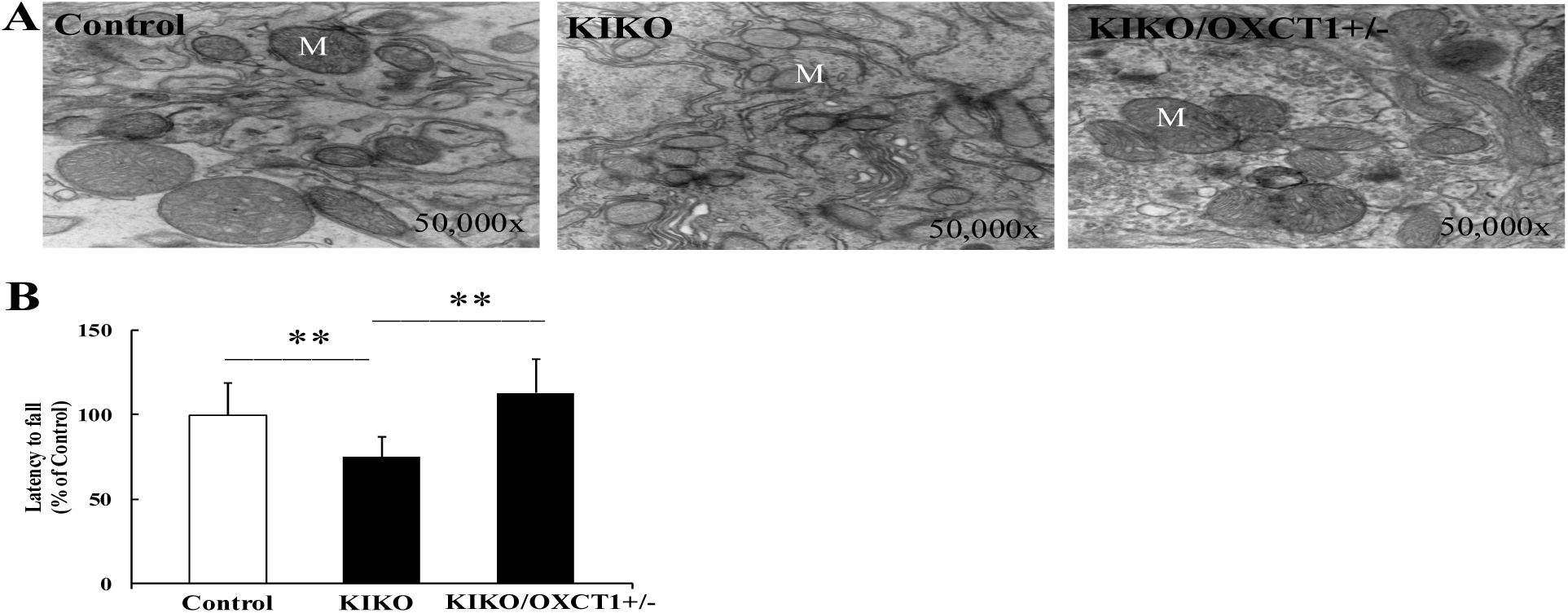
KIKO/*OXCT1*+/− mice exhibits improved mitochondrial morphology and neurobehavioral phenotype. (A) Transmission electron microscopy revealed fragmented mitochondria with reduced size in cerebellar Purkinje neurons of KIKO mice, which were normalized in KIKO/*OXCT1*⁺/⁻ mice. (B) KIKO mice demonstrated decreased latency to fall in the rotarod test, KIKO/*OXCT1*+/− mice reversed the lower latency to the levels of control mice (n=22-28). \*\**p<*0 .01. Data was shown as mean ± SE (error bars).

We next examined whether the increase in frataxin levels and mitochondrial biogenesis can reverse the neurobehavioral phenotype in KIKO/*OXCT1*+/− mice. While KIKO mice demonstrated decreased latency to fall in the rotarod test, KIKO/*OXCT1*+/− mice reversed the lower latency to the levels of control mice (**Figure 7B) (**KIKO mice: 25% decrease from control, n=22-38, *P*<0.01; KIKO/*OXCT1*+/− mice: 38% increase from KIKO mice, n=22-28, *P*<0.01), suggesting that KIKO/*OXCT1*+/− mice have improved neurobehavioral abilities.

### 3.7 Fasting increases frataxin levels and mitochondrial biogenesis markers in the cerebellum of 9-18M KIKO/*OXCT1*+/− mice

We then examined the levels of frataxin and mitochondrial biogenesis markers in 9-18M KIKO/*OXCT1*+/− mice. Consistent with our previous reports, levels of frataxin and the mitochondrial biogenesis marker Tfam were decreased at 9-18M KIKO mice (**Figure 8A and 8B) (**62% decrease for both frataxin and Tfam, n=9, *P*<0.01) as in 5-7M KIKO mice. However, unlike 5-7M KIKO/*OXCT1*+/− mice, the levels of frataxin and Tfam were unchanged in 9-18M KIKO/*OXCT1*+/− mice when compared to KIKO mice at the same age (**Figure 8A and 8B**). As increased mitochondria biogenesis is associated with ketosis in 5-7M KIKO/*OXCT1*+/− mice and fasting can induce ketosis (11), we then tested whether fasting increases mitochondria biogenesis. The 9-18M control, KIKO and KIKO/*OXCT1*+/− mice were fasted for 24 hours followed by Western blot analysis. In comparison with KIKO mice, fasting increased the levels of frataxin and the mitochondrial biogenesis marker Tfam in KIKO/*OXCT1*+/− mice (**Figure 8C and 8D) (**97% and 133% increase for frataxin and Tfam, respectively, n=6, *P*<0.01), suggesting that fasting can increase mitochondrial biogenesis and frataxin levels.

**Figure 8.**
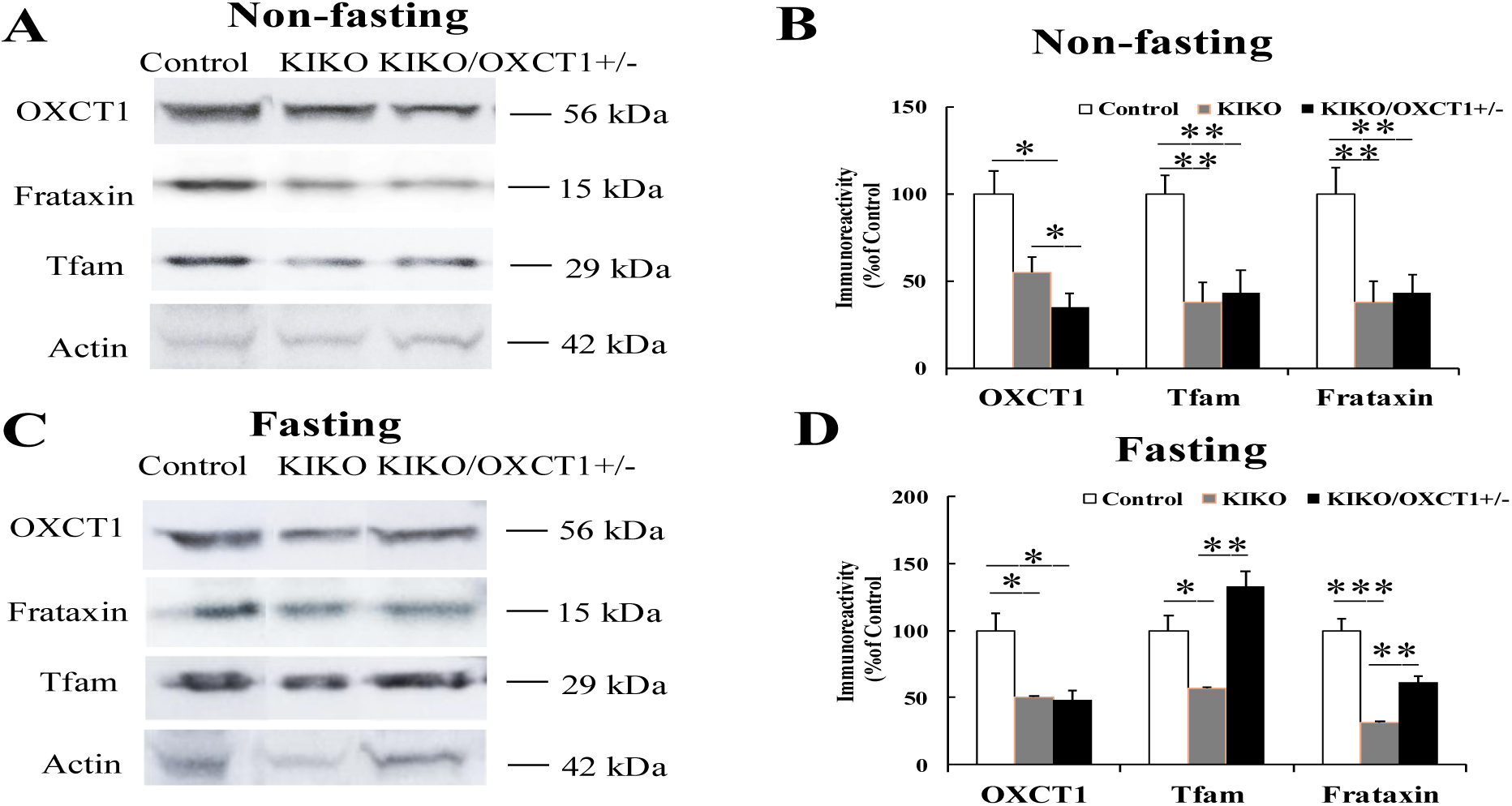
Fasting increases frataxin and mitochondrial biogenesis markers in 9-18M KIKO/*OXCT1*+/− mice. The levels of frataxin and the mitochondrial biogenesis marker Tfam were reduced in the cerebellar homogenates of 9-18M KIKO and KIKO/*OXCT1*+/− mice (A and B, n=6). Fasting partially reversed this reduction in KIKO/*OXCT1*+/− mice (C and D, n=6). * *p*<0.05, \*\**p*<0.01, \*\*\**p*<0.001. Data was shown as mean ± SE (error bars).

### 3.8 BHB treatment increase frataxin levels and mitochondrial biogenesis markers in iPSC-derived cardiomyocytes from FRDA patient

As increased frataxin levels are associated with ketosis in KIKO/*OXCT1*⁺/⁻ mice, and fasting—which induces ketosis—also elevates frataxin levels, we next treated iPSC-derived cardiomyocytes, an affected tissue in FRDA, with BHB to assess the direct effects of ketosis on frataxin expression and mitochondrial biogenesis. As shown in **Figure 9A and 9B**, iPSC-derived cardiomyocytes from FRDA patient exhibited markedly reduced frataxin levels compared with control (80% decrease, n=3, *P*<0.01). Treatment with BHB for 24 h significantly increased frataxin protein levels and Tfam expression in FRDA iPSC-derived cardiomyocytes (**Figure 9C and 9D) (**36% increase and 47% increase for frataxin and Tfam, respectively, n=6, *P*<0.05) but not in control iPSC-derived cardiomyocytes (n=6, *P*>0.05), confirming a direct effect of ketosis in promoting frataxin expression and mitochondria biogenesis.

**Figure 9.**
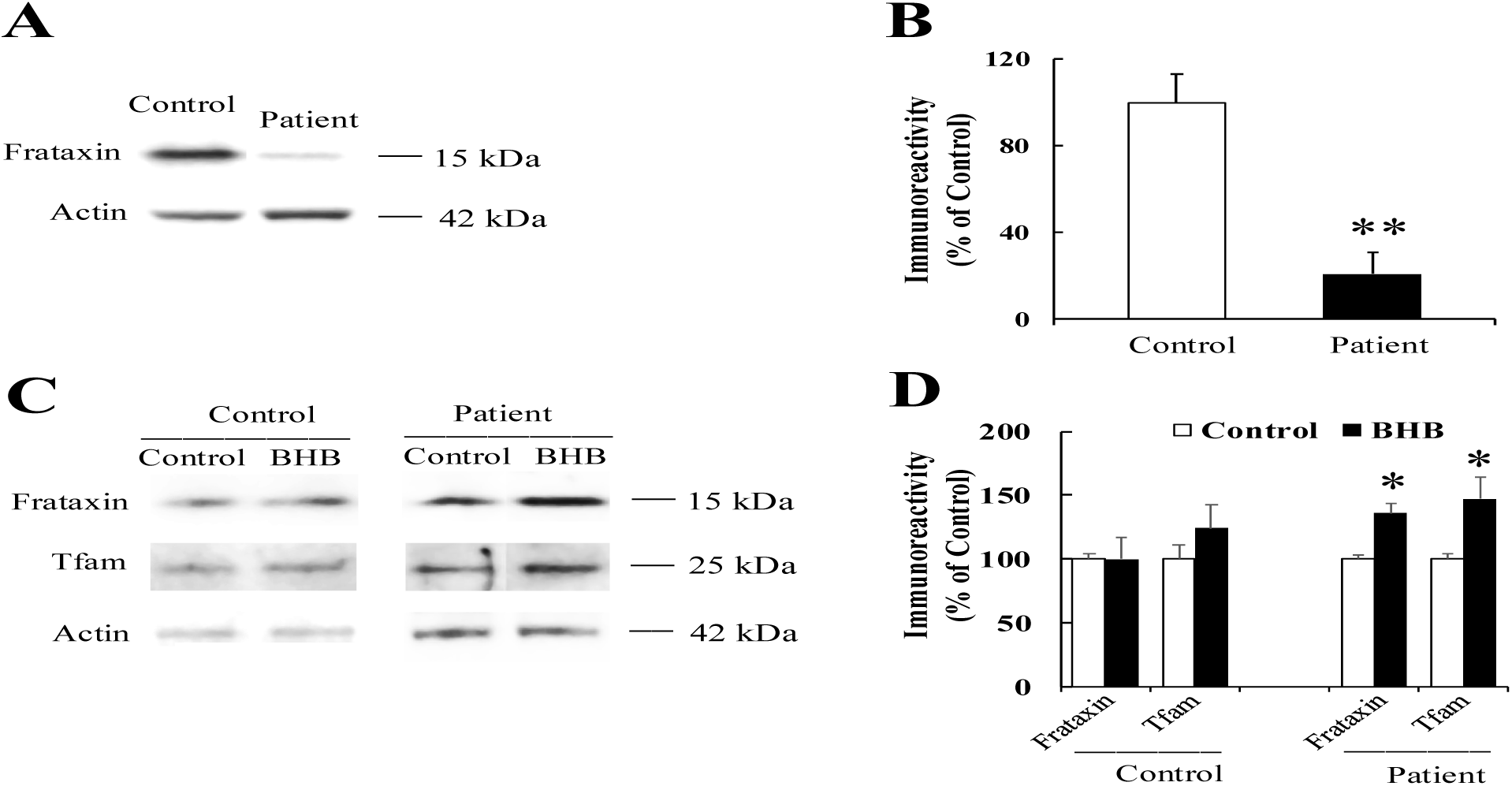
BHB treatment increases frataxin levels and mitochondrial biogenesis markers in FRDA iPSC-derived cardiomyocytes. Representative blots and bar graphs demonstrate reduced frataxin in FRDA iPSC-derived cardiomyocytes (A and B, n=3). Treatment with 10 mM BHB for 24 hours significantly increased frataxin levels and the mitochondrial biogenesis marker Tfam in FRDA iPSC-derived cardiomyocytes, but not in control iPSC-derived cardiomyocytes (C and D, n=6). * *P*<0.05, \*\**P*<0.01. Data was shown as mean ± SE (error bars).

### 3.9 Further OXCT1 deficiency causes epigenetic changes in KIKO/*OXCT1*+/− mice

The ketone body BHB is an endogenous and selective inhibitor of class I histone deacetylases (HDACs). Administration of exogenous BHB increases histone acetylation, leading to upregulation of genes involved in oxidative stress resistance and enhanced mitochondrial biogenesis in mouse tissues (35–36). To determine whether the increased mitochondrial biogenesis observed in KIKO/*OXCT1*⁺/⁻ mice is associated with ketosis-induced epigenetic regulation, we assessed the acetylation levels of histone 3 lysine 9 (H3K9) and lysine 14 (H3K14). KIKO mice exhibited significantly reduced acetylation of H3K9 and H3K14 in cerebellar homogenates compared with control mice (**Figure 10A and B**) (28% and 33% decrease for acetylated H3K9 and H3K14, respectively; n=7–8; \**P*<0.05, \*\**P*<0.01). In contrast, further OXCT1 deficiency restored H3K9 and H3K14 acetylation levels in KIKO/*OXCT1*⁺/⁻ mice to that of control mice (**Figure 10A and B**) (79% and 23% increase over KIKO mice for acetylated H3K9 and H3K14, respectively; n=7–8; \**P*<0.05, \*\**P*<0.01), suggesting that ketosis-associated epigenetic changes accompany enhanced mitochondrial biogenesis in this model.

**Figure 10.**
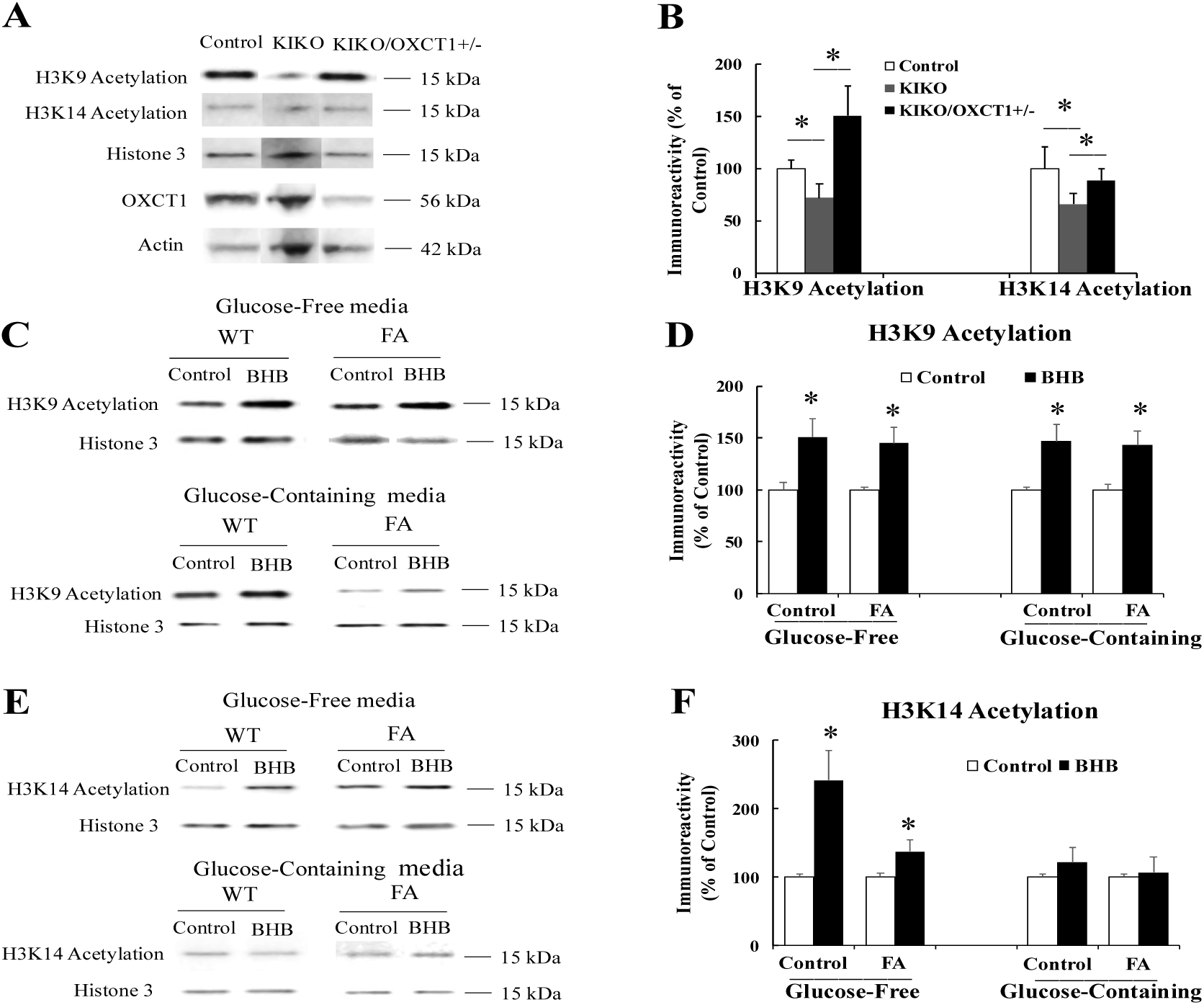
Further OXCT1 deficiency induces epigenetic changes in KIKO/*OXCT1*+/− mice. Representative blots and bar graph demonstrate reduced acetylation of H3K9 and H3K14 in the cerebellum of KIKO mice. Further OXCT1 deficiency restored H3K9 and H3K14 acetylation to normal levels (A, B) (n=7-8). Treatment with 10 mM BHB for 24 hours increased H3K9 acetylation in iPSC-derived cardiomyocytes from both control and FRDA patients under both glucose-free and glucose-containing conditions (C and D) (n=11) and increased H3K14 acetylation specifically under glucose-free condition (E and F), confirming the epigenetic effect of ketosis. * *P*<0.05. Data was shown as mean ± SE (error bars).

To further confirm that the observed epigenetic changes are directly induced by ketosis, iPSC-derived cardiomyocytes from both control and FRDA patients were treated with BHB for 24 hours under glucose-containing and glucose-free conditions. BHB treatment significantly increased H3K9 acetylation in both control and FRDA iPSC-derived cardiomyocytes under both glucose-containing and glucose-free conditions (**Figure 10C and D**) (Glucose-free condition: 51% and 45% increase for control and FRDA, respectively, n=11, *P*<0.05; Glucose-containing condition: 47% and 43% increase for control and FRDA, respectively, n=11, *P*<0.05), and increased H3K14 acetylation under glucose-free condition (**Figure 10 E and F**) (142% and 37% increase for control and FRDA, respectively, n=11, *P*<0.05). Together, these results confirm that ketosis promotes epigenetic modifications in KIKO/*OXCT1*⁺/⁻ mice through elevated H3K9 and H3K14 acetylation.

## 4 Discussion

In the present study, we demonstrate that frataxin regulates OXCT1 protein turnover by inhibiting its ubiquitination and subsequent proteasomal degradation. This effect is mediated by the N-terminal 40 amino acids of frataxin, as overexpression of this region alone is sufficient to block ubiquitin-proteasome system (UPS)-dependent OXCT1 degradation. We further establish the functional relevance of OXCT1 deficiency in both cellular and animal models of FRDA. While OXCT1 deficiency potentiates cell death *in vitro* in control and FRDA patient-derived fibroblasts. Additional OXCT1 deficiency in KIKO mice induces ketosis leading to increased frataxin protein levels and improved behavioral outcomes. The increase in frataxin levels is caused by increased mitochondria biogenesis as evidenced by upregulation of mitochondrial biogenesis markers, restoration of mitochondria size and morphology, and increased mitochondrial gene expression. These beneficial effects are supported by fasting-induced ketosis *in vivo* and by the direct effects of BHB in FRDA patient–derived iPSC cardiomyocytes. Together, these findings demonstrate a beneficial role for ketosis in restoring frataxin levels and identify ketosis-based therapies such as intermittent fasting and ketogenic diet as novel therapeutic approaches for FRDA.

The ubiquitin–proteasome system (UPS) is a central regulator of cellular protein quality control. We previously demonstrated OXCT1 undergoes proteasomal degradation and that frataxin stabilizes OXCT1 by preventing this process; however, whether OXCT1 is degraded through a ubiquitin-dependent or ubiquitin-independent proteasomal pathway remained unclear. Using cytosol-targeted and mitochondria-targeted ubiquitin constructs, we found that OXCT1 is ubiquitinated in both the cytosolic and mitochondrial compartments, with substantially greater enrichment in the mitochondrial fraction. Furthermore, inhibition of the proteasome with MG132 significantly increased levels of ubiquitinated OXCT1. Consistent with these findings, ubiquitination reduced OXCT1 protein abundance, whereas MG132 treatment restored OXCT1 levels (**Figure 1**). These results demonstrate that OXCT1 protein stability is regulated by ubiquitination. The ability of frataxin to block OXCT1 ubiquitination further underscores the molecular mechanism by which frataxin regulates OXCT1 protein turnover, thereby linking mitochondrial protein quality control to ketone body metabolism. Our findings highlight ubiquitination as a critical post-translational mechanism governing OXCT1 homeostasis and metabolic function. In further support of this model, AMPKα2 also stabilizes OXCT1 by regulating its ubiquitination (37).

Frataxin is a 210–amino acid protein critical for Fe-S cluster biogenesis (38–40). Extensive structural and biochemical studies have established that the conserved C-terminal domain of frataxin mediates binding to the NFS1–ISD11–ACP complex and the scaffold protein ISCU, facilitating controlled iron delivery for Fe–S cluster assembly while preventing iron-mediated toxicity (41–42). Consequently, the C-terminal domain has been regarded as the canonical functional region required for Fe–S cluster biogenesis and mitochondrial metabolic homeostasis, and most prior studies have focused on this region. In contrast, mapping the OXCT1-binding region of frataxin allowed us to identify a previously unrecognized functional domain within the N-terminal region of the protein. We found that the N-terminal fragment encompassing amino acids 1–40 binds OXCT1 with comparable strength to full-length frataxin and is equally effective in preventing OXCT1 degradation. Notably, GRP75, a known molecular chaperone of frataxin, also preferentially binds to the N-terminal region of frataxin (25). As OXCT1 is a downstream target of frataxin and mediates its role in ketone body metabolism, these findings suggest that the N-terminal region of frataxin not only regulates frataxin stability but also contributes to its function in ketone body metabolism and potentially other yet-unidentified pathways. Further studies examining the stability of the N-terminal fragment and identifying interaction partners mediated by this region will be necessary to fully define its functional significance.

Ketone bodies not only serve as an alternative energy source to glucose under conditions of limited glucose availability but also act as intracellular signaling molecules that exert post-translational and epigenetic effects (20, 35, 43). As a key enzyme in ketone body catabolism, the physiological consequences of OXCT1 deficiency are inherently complex, as impaired ketone utilization may also trigger adaptive metabolic responses such as ketosis. We identified a dual role for OXCT1 deficiency in FRDA. On one hand, OXCT1 deficiency potentiates cell death *in vitro* in both control and FRDA fibroblasts, consistent with previously observed defects in ATP production in the skeletal muscle cell line C2C12 (22) and with proprioceptive deficits *in vivo* (15), underscoring the critical role of ketone body utilization in maintaining cell viability and its contribution to FRDA pathology when ketone body metabolism is compromised. On the other hand, when additional OXCT1 deficiency results in ketosis, the accumulation of ketone bodies confers beneficial effects on frataxin levels and neurobehavioral outcomes, highlighting the complex and context-dependent consequences of disrupted ketone body metabolism in FRDA.

Multiple lines of evidence from our study demonstrate that ketosis increases frataxin levels, including ketosis induced by OXCT1 deficiency, fasting, and direct BHB treatment. The increase in frataxin levels is not driven by enhanced *FXN* gene transcription but instead results from ketosis-induced increased mitochondrial biogenesis, as evidenced by elevated mitochondrial biogenesis markers, increased mitochondrial gene expression, and normalization of mitochondrial size and morphology, collectively indicating improved mitochondrial homeostasis. These findings are consistent with the well-documented beneficial effects of ketosis in neurodegenerative disorders, where ketosis promotes mitochondrial biogenesis and restores mitochondrial function (21). Mitochondrial dysfunction is a central pathogenic feature of FRDA and is characterized by impaired mitochondrial biogenesis and reduced levels of multiple mitochondrial proteins involved in ATP production (33, 44). Accordingly, reversing mitochondrial dysfunction represents a key therapeutic strategy for FRDA and underlies the mechanism of action of omaveloxolone, which is currently the only approved treatment for FRDA patients aged 16 years and older. Ketosis-induced mitochondrial biogenesis not only increases frataxin levels but also elevates the expression of other mitochondrial genes, such as *ND1*, suggesting that ketosis can restore mitochondrial function through both frataxin-dependent and frataxin-independent mechanisms. The observed improvements in neurobehavioral outcomes likely reflect this restoration of mitochondrial function and further support the functional relevance of increased frataxin levels. Further studies directly assessing mitochondrial functional recovery will provide deeper insight into the therapeutic impact of ketosis.

Although the precise mechanisms linking ketosis to increased mitochondrial biogenesis in FRDA remain to be fully elucidated, our data indicate that ketosis-induced epigenetic regulation may play a key role. The ketone body BHB functions as an endogenous inhibitor of histone deacetylases (HDACs), particularly class I HDACs, thereby linking metabolic state to gene expression through epigenetic modifications (35). Administration of exogenous BHB increases global histone acetylation in mouse tissues (35). Consistent with these findings, we observed significantly increased H3K9 and H3K14 acetylation in the cerebellum of KIKO/*OXCT1*+/− mice compared with KIKO mice. Similarly, BHB treatment of iPSC-derived cardiomyocytes from both control and FRDA patients resulted in increased H3K9 and H3K14 acetylation, confirming the epigenetic effects of ketosis observed *in vivo*. These results suggest that ketosis induces epigenetic changes in KIKO/*OXCT1*+/− mice that may contribute to enhanced mitochondrial biogenesis. The significant reduction in H3K9 and H3K14 acetylation observed in the cerebellum of KIKO mice likely reflects limited acetyl-CoA availability in FRDA, which may result from both TCA cycle disruption due to impaired Fe–S cluster–containing enzymes and compromised ketone body metabolism (22, 45). Together, our findings support a role for ketosis in promoting mitochondrial biogenesis and increasing frataxin levels, highlighting ketosis-based interventions as a potential therapeutic strategy for FRDA.

## Author Contributions

Y.D. conceptualized and designed the study, conducted the experiments, supervised the project, and wrote the manuscript. M.W.A., R.S., L.V.N., J.A., O.S., J.C., S.A.H.A., C.M., P.X. performed the experiments. D.R.L. supervised the project and edited the manuscript.

## Acknowledgements

We are grateful to Dr. M. Grazia Cotticelli for assistance with qPCR and to Dr. Stewart A. Anderson and Danny Frederick for their help with iPSC culture.

